## Supplemental Table 1 for "InCellCryst - A streamlined approach to structure elucidation using *in cellulo* crystallized recombinant proteins"

**Supplementary Table 1. Overview of all vectors created for the in cellulo screening system.** Indicated are the purpose of each vector and the DNA sequence that has been cloned between the HindIII and BamHI sites of the pFastBac1 vector. Denoted are the vector-encoded amino acid sequences that are added to the N- and C-terminus of the target protein. ‘ - protease cleavage site.

| **cloning system** | **vector category** | **vector name** | **target compartment** | **N-terminal amino acid sequence** | **C-terminal amino acid sequence** | **base pair sequence between HindIII and BamHI restriction enzyme sites** |
|---|---|---|---|---|---|---|
| **first generation** | screening | pFB1 cyto | cytoplasm | MG | A | ATGGGCGCCTAA |
| | | pFB1 PTS1 | peroxisome | MG | ASKL | CGGTCCGAAGCGCGCGGAATTCATCATGGG CGCCAGCAAACTGTAA |
| | | pFB1 SS | secretory pathway | MHLMRACITFCIASTAV VAVNA‘G | A | ATGCATCTCATGCGTGCCTGCATCACATTTT GTATCGCTTCGACGGCTGTAGTCGCCGTAAA CGCCGGCGCCTAA |
| | | pFB1 SS-R | endoplasmic reticulum | MHLMRACITFCIASTAV VAVNA‘G | AKDEL | ATGCATCTCATGCGTGCCTGCATCACATTTT GTATCGCTTCGACGGCTGTAGTCGCCGTAAA CGCCGGCGCCAAAGATGAACTGTAA |
| | immunofluo rescence | pFB1 cyto HA-C | cytoplasm | MG | AYPYDVPDYA | ATGGGCGCCTACCCCTACGACGTGCCCGAC TACGCTTAA |
| | | pFB1 cyto HA-N | cytoplasm | MYPYDVPDYAG | A | ATGTACCCCTACGACGTGCCCGACTACGCT GGCGCCTAA |
| | | pFB1 SS-R HA-C | endoplasmic reticulum | MHLMRACITFCIASTAV VAVNAG | AYPYDVPDYAKD EL | ATGCATCTCATGCGTGCCTGCATCACATTTT GTATCGCTTCGACGGCTGTAGTCGCCGTAAA CGCCGGCGCCTACCCCTACGACGTGCCCGA CTACgccAAAGATGAACTGTAA |
| | | pFB1 SS-R HA-N | endoplasmic reticulum | MHLMRACITFCIASTAV VAVNAYPYDVPDYAG | AKDEL | ATGCATCTCATGCGTGCCTGCATCACATTTT GTATCGCTTCGACGGCTGTAGTCGCCGTAAA |

| | | | | | | |
|---|---|---|---|---|---|---|
| | | | | | | CGCCggcTACCCCTACGACGTGCCCGACTACGCTGGCGCCAAAGATGAACTGTAA |
| | purification | pFB1 N-His | cytoplasm | MSYYHHHHHHDYDIPTTENLYFQ‘GAMGSMG | A | ATGTCGTACTACCATCACCATCACCATCACGATTACGATATCCCAACGACCGAAAACCTGTATTTTCAGGGAGCCATGGGATCCATGGGCGCCTAA |
| | | pFB1 N-His HA-C | cytoplasm | MSYYHHHHHHDYDIPTTENLYFQ‘GAMGSMG | AYPYDVPDYA | ATGTCGTACTACCATCACCATCACCATCACGATTACGATATCCCAACGACCGAAAACCTGTATTTTCAGGGAGCCATGGGATCCATGGGCGCCTACCCCTACGACGTGCCCGACTACGCTTAA |
| **second generation** | screening | pFB1 v2 cyto | cytoplasm | MGT | AS | ATGGGTACCATTGTAGAGGATATTCCCGCCGACAGCTAGCTAA |
| | | pFB1 v2 PTS1 | peroxisome | MGT | ASKL | ATGGGTACCGAAGCTACTTAGCTGAGATTTGCCCGCTAGCAAGCTGTAA |
| | | pFB1 v2 NLS | nucleus | MGT | ASPAAKRVKLD | ATGGGTACCCGTCGTCAAGGAACAATCTTTGCTGGCTAGCCCTGCTGCTAAGAGAGTCAAGCTGGACTAA |
| | | pFB1 v2 MTS1 | mitochondrial matrix | MAARLLLRSLRVLSARSAPRPLPSARC‘SHSGT | AS | ATGGCTGCTAGACTGCTGCTGCGTTCTCTGAGAGTCCTGTCTGCTAGATCCGCTCCTAGACCTCTGCCTTCTGCTAGATGCTCTCACTCTGGTACCTATACGCTGGTGCCAGTCAAATTGCGCTAGCTAA |
| | | pFB1 v2 MTS2 | mitochondrial matrix | MATAIRLLGRRVSSWRLRPSPSPLAVPRRA‘SHSGT | AS | ATGGCTACCGCCATCAGACTGCTCGGTAGACGTGTCTCTTCCTGGAGACTCAGACCTTCTCCCTCCCCTCTGGCTGTCCCTCGTAGAGCTTCTCACTCTGGTACCACTTCGTCGCATTGTGTAGGCCAAAGCTAGCTAA |
| | | pFB1 v2 SS | secretory pathway | MHLMRACITFCIASTAVVAVNA‘GT | AS | ATGCACCTGATGAGAGCTTGCATCACCTTCTGCATCGCTTCTACCGCTGTCGTCGCTGTCAA |

| | | | | | | |
|---|---|---|---|---|---|---|
| | | | | | | CGCTGGTACCTCCCCTATGAACGATGTGTCGTGAAGCTAGCTAA |
| | | pFB1 v2 SS-R | endoplasmic reticulum | MHLMRACITFCIASTAVVAVNA‘GT | ASKDEL | ATGCACCTGATGAGAGCTTGCATCACCTTCTGCATCGCTTCTACCGCTGTCGTCGCTGTCAACGCTGGTACCTTCGATGGCTCTAATTGCAACAGGGGCTAGCAAGGACGAACTGTAA |
| | immunofluorescence | pFB1 v2 cyto HA-N | cytoplasm | MYPYDVPDYAGT | AS | ATGTACCCTTACGACGTGCCTGACTACGCTGGTACCGTTTCTTACGCAGGCCACGTAAGTAGCTAGCTAA |
| | | pFB1 v2 cyto HA-C | cytoplasm | MGT | ASYPYDVPDYA | ATGGGTACCTATGCTTGGAGTCCTAACCCAGTAGGCTAGCTACCCTTACGACGTGCCTGACTACGCCTAA |
| | | pFB1 v2 PTS1 HA-C | peroxisome | MGT | ASYPYDVPDYASKL | ATGGGTACCCAGACTTATGCGTGGTTACGACTCAGCTAGCTACCCTTACGACGTGCCTGACTACGCTTCTAAGCTGTAA |
| | | pFB1 v2 NLS HA-C | nucleus | MGT | ASYPYDVPDYAPAAKRVKLD | ATGGGTACCCGAGATAGGATCTCGCTTCACGTTAGCTAGCTACCCTTACGACGTGCCTGACTACGCTCCTGCTGCTAAGAGAGTCAAGCTGGACTAA |
| | | pFB1 v2 MTS1 HA-C | mitochondrial matrix | MAARLLLRSLRVLSARSAPRPLPSARC‘SHSGT | ASYPYDVPDYA | ATGGCTGCTAGACTGCTGCTGCGTTCTCTGAGAGTCCTGTCTGCTAGATCCGCTCCTAGACCTCTGCCTTCTGCTAGATGCTCTCACTCTGGTACCGTAACGTTGGGTTCTCTAAGCGTCCGCTAGCTACCCTTACGACGTGCCTGACTACGCCTAA |
| | | pFB1 v2 MTS2 HA-C | mitochondrial matrix | MATAIRLLGRRVSSWRLRPSPSPLAVPRRA‘SHSGT | ASYPYDVPDYA | ATGGCTACCGCCATCAGACTGCTCGGTAGACGTGTCTCTTCCTGGAGACTCAGACCTTCTCCCTCCCCTCTGGCTGTCCCTCGTAGAGCTTCTCACTCTGGTACCGCCTCAAGGAGAGTTTGGTCCTAATGCTAGCTACCCTTACGACGTGCCTGACTACGCCTAA |

| | | | | | | |
|---|---|---|---|---|---|---|
| | | pFB1 v2 SS-R HA-C | endoplasmic reticulum | MKLSLVAAML LLLSAARA‘GT | ASYPYDVPDYAKDEL | ATGCACCTGATGAGAGCTTGCATCACCTTCTGCATCGCTTCTACCGCTGTCGTCGCTGTCAACGCTGGTACCTATCCGTAAGTTCGTAAGCACCTGGGCTAGCTACCCTTACGACGTGCCTGACTACGCTAAGGACGAACTGTAA |
| | | pFB1 v2 SS-R HA-N | endoplasmic reticulum | MKLSLVAAML LLLSAARA‘YPYDVPDYAGT | ASKDEL | ATGCACCTGATGAGAGCTTGCATCACCTTCTGCATCGCTTCTACCGCTGTCGTCGCTGTCAACGCTTACCCTTACGACGTGCCTGACTACGCTGGTACCATCGGGAGATTCGCAAACGTTTCTCGCTAGCAAGGACGAACTGTAA |
| | fluores-cence | pFB1 v2 cyto mTurq2-C | cytoplasm | GT | ASVSKGEELFTGVVPILVELDGDVNGHKFSVSGEGEGDATYGKLTLKFICTTGKLPVPWPTLVTTLSWGVQCFARYPDHMKQHDFFKSAMPEGYVQERTIFFKDDGNYKTRAEVKFEGDTLVNRIELKGIDFKEDGNILGHKLEYNYFSDNVYITADKQKNGIKANFKIRHNIEDGGVQLADHYQQNTPIGDGPVLLPDNHYLSTQSKLSKDPNEKRDHMVLLEFVTAAGITLGMDELYK | ATGGGTACCATTGTAGAGGATATTCCCGCCGACAGCTAGCGTGAGCAAGGGCGAGGAGCTGTTCACCGGGGTGGTGCCCATCCTGGTCGAGCTGGACGGCGACGTAAACGGCCACAAGTTCAGCGTGTCCGGCGAGGGCGAGGGCGATGCCACCTACGGCAAGCTGACCCTGAAGTTCATCTGCACCACCGGCAAGCTGCCCGTGCCCTGGCCCACCCTCGTGACCACCCTGTCCTGGGGCGTGCAGTGCTTCGCCCGCTACCCCGACCACATGAAGCAGCACGACTTCTTCAAGTCCGCCATGCCCGAAGGCTACGTCCAGGAGCGCACCATCTTCTTCAAGGACGACGGCAACTACAAGACCCGCGCCGAGGTGAAGTTCGAGGGCGACACCCTGGTGAACCGCATCGAGCTGAAGGGCATCGACTTCAAGGAGGACGGCAACATCCTGGGGCACAAGCTGGAGTACAACTACTTTAGCGACAACGTCTATATCACCGCCGACAAGCAGAAGAACGGCATCAAGGCCAACTTCAAGATCCGCCACAACATCGAGGACGGCGGCGTGCAGCTCGCCGACCACTACCAGCAGAACACCCCCATCGGCGACGGCCCCGTGCTGCTGCCCGACAACCACTACCTGAGCACCCAGTCCAAGCTGAGCAAAGACCCCAACGAGAAGCGCGATCACATGGTCCTGCTGGAGTTCGTGACCGCCGCCGGGATCACTCTCGGCATGGACGAGCTGTACAAGTAA |

| | | | | | | |
|---|---|---|---|---|---|---|
| | | pFB1 v2 MTS1 mTurq2-C | mitochondrial matrix | MAARLLLRSLRVLSARSAPRPLPSARC‘SHSGT | ASVSKGEELFTGVVPILVELDGDVNGHKFSVSGEGEGDATYGKLTLKFICTTGKLPVPWPTLVTTLSWGVQCFARYPDHMKQHDFFKSAMPEGYVQERTIFFKDDGNYKTRAEVKFEGDTLVNRIELKGIDFKEDGNILGHKLEYNYFSDNVYITADKQKNGIKANFKIRHNIEDGGVQLADHYQQNTPIGDGPVLLPDNHYLSTQSKLSKDPNEKRDHMVLLEFVTAAGITLGMDELYK | ATGGCTGCTAGACTGCTGCTGCGTTCTCTGAGAGTCCTGTCTGCTAGATCCGCTCCTAGACCTCTGCCTTCTGCTAGATGCTCTCACTCTGGTACCGTAACGTTGGGTTCTCTAAGCGTCCGCTAGCGTGAGCAAGGGCGAGGAGCTGTTCACCGGGGTGGTGCCCATCCTGGTCGAGCTGGACGGCGACGTAAACGGCCACAAGTTCAGCGTGTCCGGCGAGGGCGAGGGCGATGCCACCTACGGCAAGCTGACCCTGAAGTTCATCTGCACCACCGGCAAGCTGCCCGTGCCCTGGCCCACCCTCGTGACCACCCTGTCCTGGGGCGTGCAGTGCTTCGCCCGCTACCCCGACCACATGAAGCAGCACGACTTCTTCAAGTCCGCCATGCCCGAAGGCTACGTCCAGGAGCGCACCATCTTCTTCAAGGACGACGGCAACTACAAGACCCGCGCCGAGGTGAAGTTCGAGGGCGACACCCTGGTGAACCGCATCGAGCTGAAGGGCATCGACTTCAAGGAGGACGGCAACATCCTGGGGCACAAGCTGGAGTACAACTACTTTAGCGACAACGTCTATATCACCGCCGACAAGCAGAAGAACGGCATCAAGGCCAACTTCAAGATCCGCCACAACATCGAGGACGGCGGCGTGCAGCTCGCCGACCACTACCAGCAGAACACCCCCATCGGCGACGGCCCCGTGCTGCTGCCCGACAACCACTACCTGAGCACCCAGTCCAAGCTGAGCAAAGACCCCAACGAGAAGCGCGATCACATGGTCCTGCTGGAGTTCGTGACCGCCGCCGGGATCACTCTCGGCATGGACGAGCTGTACAAGTAA |
| | | pFB1 v2 MTS2 mTurq2-C | mitochondrial matrix | MATAIRLLGRRVSSWRLRPSPSPLAVPRRA‘SHSGT | ASVSKGEELFTGVVPILVELDGDVNGHKFSVSGEGEGDATYGKLTLKFICTTGKLPVPWPTLVTTLSWGVQCFARYPDHMKQHDFFKSAMPEGYVQERTIFFKDDGNY | ATGGCTACCGCCATCAGACTGCTCGGTAGACGTGTCTCTTCCTGGAGACTCAGACCTTCTCCCTCCCCTCTGGCTGTCCCTCGTAGAGCTTCTCACTCTGGTACCGCCTCAAGGAGAGTTTGGTCCTAATGCTAGCGTGAGCAAGGGCGAGGAGCTGTTCACCGGGGTGGTGCCCATCCTGGTCGAGCTGGACGGCGACGTAAACGGCCACAAGTTCAGCGTGTCCGGCGAGGGCGAGGGCGATGCCACCTACGGCAAGCTGACCCTGAAGTT |

| | | | | | | |
|---|---|---|---|---|---|---|
| | | | | | KTRAEVKFEGDT<br>LVNRIELKGIDFK<br>EDGNILGHKLEY<br>NYFSDNVYITAD<br>KQKNGIKANFKIR<br>HNIEDGGVQLAD<br>HYQQNTPIGDGP<br>VLLPDNHYLSTQ<br>SKLSKDPNEKRD<br>HMVLLEFVTAAGI<br>TLGMDELYK | CATCTGCACCACCGGCAAGCTGCCCGTGCC<br>CTGGCCCACCCTCGTGACCACCCTGTCCTG<br>GGGCGTGCAGTGCTTCGCCCGCTACCCCGA<br>CCACATGAAGCAGCACGACTTCTTCAAGTCC<br>GCCATGCCCGAAGGCTACGTCCAGGAGCGC<br>ACCATCTTCTTCAAGGACGACGGCAACTACA<br>AGACCCGCGCCGAGGTGAAGTTCGAGGGC<br>GACACCCTGGTGAACCGCATCGAGCTGAAG<br>GGCATCGACTTCAAGGAGGACGGCAACATC<br>CTGGGGCACAAGCTGGAGTACAACTACTTTA<br>GCGACAACGTCTATATCACCGCCGACAAGC<br>AGAAGAACGGCATCAAGGCCAACTTCAAGAT<br>CCGCCACAACATCGAGGACGGCGGCGTGCA<br>GCTCGCCGACCACTACCAGCAGAACACCCC<br>CATCGGCGACGGCCCCGTGCTGCTGCCCGA<br>CAACCACTACCTGAGCACCCAGTCCAAGCT<br>GAGCAAAGACCCCAACGAGAAGCGCGATCA<br>CATGGTCCTGCTGGAGTTCGTGACCGCCGC<br>CGGGATCACTCTCGGCATGGACGAGCTGTA<br>CAAGTAA |
| | purification | pFB1 v2 N-His | cytoplasm | MSYYHHHHHHDYDYPYD<br>VPDYAIPTTENLYFQ‘SGT | AS | ATGTCTTACTACCACCATCACCACCATCACG<br>ACTACGACTACCCTTACGACGTGCCTGACTA<br>CGCTATCCCTACCACCGAAAACCTGTACTTC<br>CAGTCTGGTACCTTGTCTAAGCCGAACGTAT<br>CGCATGGCTAGCTAA |
| | | pFB1 v2 SS N-His | secretory pathway | MKLSLVAAML<br>LLLSAARA‘SYYHHHHHHD<br>YDYPYDVPDYAIPTTENLY<br>FQ‘SGT | AS | ATGCACCTGATGAGAGCTTGCATCACCTTCT<br>GCATCGCTTCTACCGCTGTCGTCGCTGTCAA<br>CGCTTCTTACTACCACCATCACCACCATCAC<br>GACTACGACTACCCTTACGACGTGCCTGACT<br>ACGCTATCCCTACCACCGAAAACCTGTACTT<br>CCAGTCTGGTACCGCGTTCTAAATGTACCCG<br>TCAGGATGCTAGCTAA |
