## Supplemental Table 2 for "InCellCryst - A streamlined approach to structure elucidation using *in cellulo* crystallized recombinant proteins"

**Supplementary Table 2: Diffraction data processing summary.**

| **Dataset** | **HEX-1 cyto** | **HEX-1 cyto** | **HEX-1 ori** | **HEX-1 ori** | **HEX-1 cyto v2** | **IMPDH ori** | **IMPDH ori** | **IMPDH cyto** |
|---|---|---|---|---|---|---|---|---|
| Data collection date | 2017.07.02 | 2021.08.24 | 2019.11.26 | 2019.11.26 | 2021.08.24 | 2019.11.26 | 2019.11.26 | 2019.09.17 |
| Sample temperature [K] | 296 | 100 | 100 | 100 | 100 | 100 | 100 | 100 |
| Processing software | CrystFEL | CrystFEL | CrystFEL | XDS | CrystFEL | CrystFEL | XDS | CrystFEL |
| PDB code | 8C5K | 8CD4 | 8CD5 | 8CGX | 8CD6 | 8C53 | 8CGY | 8C51 |
| Spacegroup | P65 2 2 | P65 2 2 | P65 2 2 | P65 2 2 | P65 2 2 | P4 21 2 | P4 21 2 | P4 21 2 |
| Unit cell | 58.65 58.65 191.46 90 90 120 | 57.39 57.39 189.96 90 90 120 | 57.23 57.23 198.15 90 90 120 | 57.24 57.24 198.19 90 90 120 | 56.7 56.7 196.8 90 90 120 | 207.12 207.12 92.54 90 90 90 | 207.1 207.1 92.5 90 90 90 | 205.77 205.77 92.08 90 90 90 |
| Resolution Range | 49.09 - 2.16 (2.237 - 2.16) | 50.19 - 1.83 (1.895 - 1.83) | 48.08 - 1.56 (1.616 - 1.56) | 33.03 - 1.85 (1.916 - 1.85) | 47.64 - 1.85 (1.916 - 1.85) | 57.38 – 2.30 (2.33 - 2.299) | 92.62 - 3.0 (3.107 - 3.0) | 77.82 – 2.40 (2.456 - 2.399) |
| Total Reflections | 2,954,673 | 24,754,589 | 210,384,917 | 80,449,866 | 4,735,984 | 326,687,476 | 97,508,239 | 139,358,146 |
| Unique Reflections | 11,210 (1,066) | 17.908 | 28.553 | 17,305 (1,674) | 17 | 89,070 (5,857) | 40,798 (9,383) | 77,541 (5,119) |
| Multiplicity | 246 (68) | 1,382 (100) | 7,367 (610) | 4,639 (4,197) | 278 (115.9) | 3,667 (1,246) | 474 (2,177) | 1,797 (1,018) |
| Completeness (%) | 99.96 (99.72) | 99.92 (99.31) | 99.83 (98.82) | 99.8 (98.3) | 99.81 (99.09) | 99.88 (99.09) | 99.8 (99.2) | 99.92 (99.32) |
| SNR/ < I/σ(I)> | 10.41 (0.90) | 22.16 (0.64) | 31.47 (0.59) | 35.15 (3.30) | 8.15 (0.53) | 10.06 (0.58) | 20.37 (4.21) | 9.13 (0.89) |
| Wilson B-factor | 24,10 | 31,70 | 27,56 | 31,60 | 34,14 | 53.06 | 53.76 | 61,37 |
| R-meas | - | - | - | 117.0 (56,232.3) | - | - | 341.1 (8,831.4) | - |
| R-split | 5.86 (87.1) | 3.40 (168.07) | 1.95 (188.97) | - | 7.95 (184.98) | 8.62 (199.52) | - | 8.90 (180.99) |
| CC1/2 | 0.998 (0.595) | 0.9994(0.2745) | 0.9997 (0.2865) | 0.999(0.939) | 0.9953 (0.4226) | 0.9976 (0.1957) | 0.981 (0.870) | 0.9973 (0.2331) |
| CC* | 0.999 (0.864) | 0.9998(0.6564) | 0.9999 (0.6674) | - | 0.9988 (0.7707) | 0.9994 (0.5722) | - | 0.9993 (0.6149) |
| **Refinement** | | | | | | | | |
| No. collected images | 595,419 | 119,053 | 56,891 | 56,891 | 114,498 | 96,949 | 96,949 | 82,534 |
| hits / lattices | 91,354/ 57,052 | 44,200/41,591 | 54,077/131,442 | 7,922 ind. Crystals (each up to 7 cons. frames) after filtering | 24,520/21,835 | 75,741/82,831 | 6,001 ind. Crystals (each up to 11 cons. frames) after filtering | 55,854/71,502 |
| lattices after stream_grep | 38,394 | - | - | - | 6,117 | - | - | - |
| removed salt intensities | - | 0.02 % | 0.03 % | - | 0.01 % | 7.6 % | - | 3.8 % |
| Reflections used in refinement | 11,207 (1,066) | 17,807 (1,718) | 28,419 (2,729) | 18,624 (1,699) | 16,899 (1,631) | 88932 (8710) | 40,788 (3982) | 73698 (7591) |
| Reflections used for R-free | 1,120 (106) | 1,323 (127) | 1,473 (139) | 1,268 (117) | 1,045 (102) | 1078 (107) | 1,598 (156) | 1058 (109) |
| R-work | 0.182 (0.320) | 0.1923 (0.3548) | 0.1993 (0.4334) | 0.2061 (0.3500) | 0.2034 (0.3964) | 0.2132 (0.3772) | 0.2014 (0.3415) | 0.2097 (0.3853) |
| R-free | 0.231 (0.362) | 0.2148 (0.3489) | 0.2159 (0.4060) | 0.2352 (0.4580) | 0.2369 (0.4258) | 0.2431 (0.3386) | 0.2388 (0.4156) | 0.2372 (0.4257) |
| Number of non-hydrogen atoms | 1.123 | 1.185 | 1.226 | 1202 | 1.174 | 7.081 | 6.852 | 6.887 |
| macromolecules | 1,100 | 1.095 | 1.128 | 1.124 | 1.090 | 6.799 | 6.716 | 6.791 |
| ligands | 0 | 0 | 0 | 0 | 0 | 182 | 182 | 182 |
| solvent | 23 | 90 | 98 | 78 | 84 | 148 | 2 | 72 |
| Protein residues | 141 | 143 | 147 | 148 | 143 | 894 | 894 | 896 |
| RMS bonds (Å) | 0,016 | 0,011 | 0,004 | 0,006 | 0,008 | 0,004 | 0,005 | 0,005 |
| RMS angles (°) | 1.26 | 1,02 | 0,71 | 0,86 | 0,91 | 0,62 | 0,63 | 0,67 |
| Ramachandran favored (%) | 96.4 | 97,87 | 98,62 | 97,95 | 98,58 | 97,4 | 95,46 | 96,04 |
| Ramachandran allowed (%) | 44715 | 2,13 | 0,69 | 1,37 | 1,42 | 2,6 | 4,54 | 3,85 |
| Ramachandran outliers (%) | 0 | 0 | 0,69 | 0,68 | 0 | 0 | 0 | 0,11 |
| Rotamer outliers (%) | 0 | 0 | 0 | 0 | 0 | 0,97 | 0,14 | 0,83 |
| Clash score | 3.15 | 3,65 | 4,44 | 4,03 | 6,89 | 5,26 | 12,78 | 6,15 |
| Average B-factor | 57.79 | 39,19 | 41,25 | 42,06 | 40,88 | 59,98 | 52,12 | 68,55 |
| macromolecules | 57.94 | 38,64 | 40,6 | 41,65 | 40,27 | 59,87 | 51,77 | 68,29 |
| ligands | - | - | - | - | - | 61,64 | 58,01 | 75,04 |
| solvent | 50.71 | 45,94 | 48,67 | 47,91 | 48,73 | 57,22 | 34,41 | 64,03 |
| TLS refinement groups | - | 2 | 8 | 3 | - | - | 11 | 12 |
| mean RMSD to ref. structure (6rfu f. IMPDH; 7asx f. HEX-1) | 0,4990 | 0,2600 | 0,4240 | 0,4226 | 0,5203 | 0,3650 | 0,4143 | 0,4150 |
| max RMSD to ref. structure (6rfu f. IMPDH; 7asx f. HEX-1) | 2,7690 | 0,6520 | 2,1740 | 2,1700 | 2,0840 | 7,0810 | 7,0310 | 7,0040 |
