## Supplemental Table 3 for "InCellCryst - A streamlined approach to structure elucidation using *in cellulo* crystallized recombinant proteins"

**Supplementary Table 3. Overview of all cloning primers used.** Indicated are

| Name | Sequence (5‘ → 3‘) |
|---|---|
| **CatB fwd** | GATCGGATCCATGCATCTCATGCGTGCCT |
| **CatB KDEL rev** | GATCCTCGAGCTACAGCTCATCCTTCGCCGTGTTGGG |
| **IMPDH v1 fwd** | GAAAACACCAACCTACGCACCA |
| **IMPDH v1 rev** | GGCAAAGAGTTTCCTCTCGTAGTGG |
| **IMPDH QC fwd** | GCGCTGGCGGTTGGAGCCAACGTGGCGATG |
| **IMPDH QC rev** | CATCGCCACGTTGGCTCCAACCGCCAGCGC |
| **EGFP-µNS v2 fwd** | GATCGGTACCATGGTGAGCAAGGGC |
| **EGFP-µNS v2 rev** | GATCGCTAGCCAATCGTACGTTAGCGGAACG |
| **HA C-term check rev** | CATCTTTGGCGTAGTCGGG |
| **HA C-term cyto fwd** | GATCGGATCCATGGGCGCCTACCCCTACGACGTGCCCGA |
| **HA C-term cyto rev** | GATCAAGCTTTTAAGCGTAGTCGGGCACGTCGTAGGGGT |
| **HA C-term fwd** | GCCTACCCCTACGACGTGCCCGACTAC |
| **HA C-term rev** | GTAGTCGGGCACGTCGTAGGGGTAGGC |
| **HA N-term check rev** | CTTTGGCGCCAGCGTAG |
| **HA N-term cyto fwd** | GATCGGATCCATGTACCCCTACGACGTGCCCGACTACGC |
| **HA N-term cyto rev** | GATCAAGCTTTTAGGCGCCAGCGTAGTCGGGCACGTCGT |
| **HA N-term fwd** | TACCCCTACGACGTGCCCGACTACGCTGGC |
| **HA N-term rev** | GCCAGCGTAGTCGGGCACGTCGTAGGGGTA |
| **pFB1 fwd Seq v1** | TAAAATGATAACCATCTCGC |
| **pFB1 fwd Seq v2** | TTCATACCGTCCCACCATCG |
| **pFB1 fwd Seq v3** | GTTGGCTACGTATACTCCGGA |
| **pFB1 rev Seq** | TTCAGGTTCAGGGGGAGGTG |
| **pFB1 rev Seq v2** | ACAAACCACAACTAGAATGCAGTG |
| **luciferase v1 fwd** | GAAGACGCCAAAAACATAAAGAA |
| **luciferase$^{-}$ v1 rev** | CTTTCCGCCCTTCTTGGC |
| **luciferase$^{+}$ v1 rev** | CAATTTGGACTTTCCGCCCTTC |
| **luciferase QC fwd** | GAAAGGCCCGGCTCCATTCTATCCTC |
| **luciferase QC rev** | GAGGATAGAATGGAGCCGGGCCTTTC |
| **HEX-1 v1 fwd** | TACTACGACGACGACGCTCACG |
| **HEX-1 ori fwd** | CTAGGGATCCATGGGCTACTACGACGACGAC |
| **HEX-1 ori rev** | CTAGAAGCTTTTAGAGGCGGGAACCGTGG |
| **HEX-1 v1 rev** | GAGGCGGGAACCGTGGACG |
| **HEX-1 v2 fwd** | CTAGGGTACCGGCTACTACGACGACGACG |
| **HEX-1 v2 rev** | CTAGGCTAGCGAGGCGGGAACCGTGG |
| **pUC/M13 fwd** | CCCAGTCACGACGTTGTAAAACG |
| **pUC/M13 rev** | AGCGGATAACAATTTCACACAGG |
| **FP v1 fwd** | GTGAGCAAGGGCGAGGAG |
| **FP v1 rev** | CTTGTACAGCTCGTCCATGCCG |
| **mTurq v2 fwd** | GATCGCTAGCGTGAGCAAGGGCGAG |
| **mTurq v2 rev** | GATCAAGCTTACTTGTACAGCTCGTCC |
